# Waddington Revisited: Organ transformation by environmental disruption of epigenetic memory

**DOI:** 10.1101/482216

**Authors:** Orli Snir, Michael Elgart, Filippo Ciabrelli, Shlomi Dagan, Iris Aviezer, Elizabeth Stoops, Giacomo Cavalli, Yoav Soen

## Abstract

Despite major progress in mechanistic understanding of epigenetic reprogramming of cells, the basis of ‘organ reprograming’ by (epi-)gene-environment interactions remained largely obscured. Here we use the ether-induced haltere-to-wing transformations as a model for epigenetic “reprogramming” at the whole organism level. Our findings support a mechanistic chain of events explaining why and how brief embryonic exposure to ether leads to organ transformation manifested at the larval stage and on. We show that ether interferes with protein integrity in the egg leading to altered deployment of Hsp90 and repression of Trithorax-mediated establishment of H3K4 tri-methylations. This repression pre-disposes early methylated Ubx targets and wing genes for later up-regulation in the larval haltere disc, hence the wing-like outcome. Consistent with compromised protein integrity during the exposure, the severity of bithorax transformation is increased by genetic or chemical reduction of Hsp90 function. Moreover, a joint reduction in *Hsp90* and *trx* gene dosage can cause bithorax transformations without exposure to ether. These findings implicate environmental disruption of protein integrity at the onset of histone methylations with a modification of epigenetic memory, which in turn, supports a morphogenetic shift towards an ancestral-like body plan. The morphogenetic impact of chaperone response during a major setup of epigenetic patterns may be a general scheme for organ reprogramming by environmental cues.

## Introduction

Determination of cell and tissue identities in flies are established during embryonic development and maintained by epigenetic means, particularly by the Polycomb and Trithorax systems^1,2^. Early embryonic exposure to environmental stimuli (e.g. ether vapor and heat) can interfere with these identities and induce homeotic transformations, such as haltere-to-wing (bithorax) phenocopies^3–7^. These phenocopies can be further stabilized (assimilated) by repeated exposures and selections over several generations^6,8,9^. This provided a striking example for environmental induction of a gross morphogenetic transformation that can be rapidly stabilized, but the mechanistic basis of this phenomenon remained largely unknown. While the similarity to mutation phenotypes of homeotic genes (e.g. *Ubx* and *trx*) suggests their potential involvement in the induced transformation^9,10^, it is not clear how these (or other) genes mediate such a reaction to brief exposure to ether. Moreover, the epigenetic basis of the induced haltere-to-wing transformations has not been yet laid out.

By analyzing the effects of exposure to ether, we provide evidence for a mechanistic chain of events, connecting early exposure to ether with induction of bithorax phenocopies. We show that ether vapor disrupts native folding of proteins in the embryo and severely compromises H3K4 tri-methylation by trx at a critical stage of development. We found that this disruption of active chromatin marks is less pronounced in actively expressed genes such as wing development genes. The preferential retention of H3K4me3 marks pre-disposes these wing genes for later up-regulation in the larval haltere disc. In support of the causal contribution of interaction between proteotoxic stress and reduced H3K4 tri-methylations, we found that chemical or genetic reduction of Hsp90 enhances the bithorax reaction and that the induction is further aggravated by joint reduction in *Hsp90* and *trx* gene dosage (in *Hsp83*^e6A^/*trx*^1^ double heterozygotes). Consistent with a reported dependence of trx on Hsp90 function^11^, joint reduction in *Hsp90* and *trx* gene dosage also recapitulated the spontaneous generation of bithorax transformations that were observed in *trx*^1-/-^ homozygotes, but not in *trx*^1^ heterozygotes. Altogether, these findings link the proteotoxic stress of ether to excess demand for Hsp90 function which interferes with initial epigenetic patterning and predisposes wing genes for later up-regulation in the haltere disc. This chain of events can also account for induction of bithorax phenocopies in response to brief exposure to heat at this critical stage in development^12,13^.

## Results

### Ether suppresses H3K4me3 tri-methylation in the early embryo

We established effective conditions for bithorax induction in response to short (30min) exposure of early stage (3hr old) *yw* embryos to ether vapor (Fig. 1A). To quantify the reaction, we scored the number and fraction of adult flies (or failed-to-eclose pupae) exhibiting a bithorax phenocopy, defined as any morphological abnormality in the 3rd thoracic segment that increases resemblance to the 2nd thoracic segment^3,6,7,14^ (Fig. 1B). Bithorax phenocopy was induced in ∼9% of the individuals without a significant impact on the number of pupae (Fig. 1C, D). To investigate how embryonic exposure leads to homeotic transformations at a substantially later stage, we analyzed histone methylation and transcriptional changes shortly after exposure (Fig. 1A). Active and repressive chromatin marks were analyzed by ChIP-seq with antibodies specific for Histone 3 tri-methylation of Lysine 4 (H3K4me3) and Lysine 27 (H3K27me3), respectively^15^. Comparison between methylation states in different samples was preceded by global percentile normalization of read counts applied to 100bp genomic segments of each sample^16^. Differences between normalized counts of exposed (‘Ether’) vs. non-exposed embryos (‘Ctrl’) revealed genome-wide (albeit not uniform) decrease in regions with H3K4 tri-methylation, without any noticeable change in tri-methylation of H3K27 (Fig. 1E vs. Fig. 1F; Supplementary Fig. S1, S2A). Predominant suppression of H3K4me3 was also reflected by site-specific ratio of H3K4me3 to H3K27me3, determined by integrating normalized counts over each gene region and dividing the H3K4me3 level by the level of H3K27me3 at the same region. In ether-exposed vs. control embryos, the H3K4me3/H3K27me3 ratio was significantly lower in 88% of the genes and higher in only 5% of the genes (Fig. 1G). Unlike the extensive suppression of H3K4me3, the immediate transcriptional response was very mild (Fig. 1H; Supplementary Fig. S2B,C; Supplementary Spreadsheet S1). Moreover, the set of differentially expressed genes was not enriched with wing genes (Fig. 1I), suggesting that the induced predisposition towards wing might be primarily specified by site-specific changes in histone methylation at the time of exposure. This was supported by significantly higher abundance of H3K4 tri-methylations at loci of wing genes (Fig. 1J ; Supplementary Fig. S2D, E) as well as by enrichment of wing genes in loci that exhibited the highest retention of H3K4 tri-methylation following exposure to ether (Fig. 1K; Supplementary Table S1). High retention of K4 tri-methylation was also correlated with high levels of expression at around the time of exposure (Fig. 1L, Supplementary Fig. S2F-I), suggesting that high expression of wing genes during the exposure, contributes to preferential retention of K4 tri-methylation in these genes. Given the reported contribution of H3K4 tri-methylation to long-term stability of active state of transcription^17,18^, higher abundance of H3K4 tri-methylation at wing genes could favor sustained expression of these genes at higher levels compared to the respective levels in non-exposed embryos.

**Figure 1.**
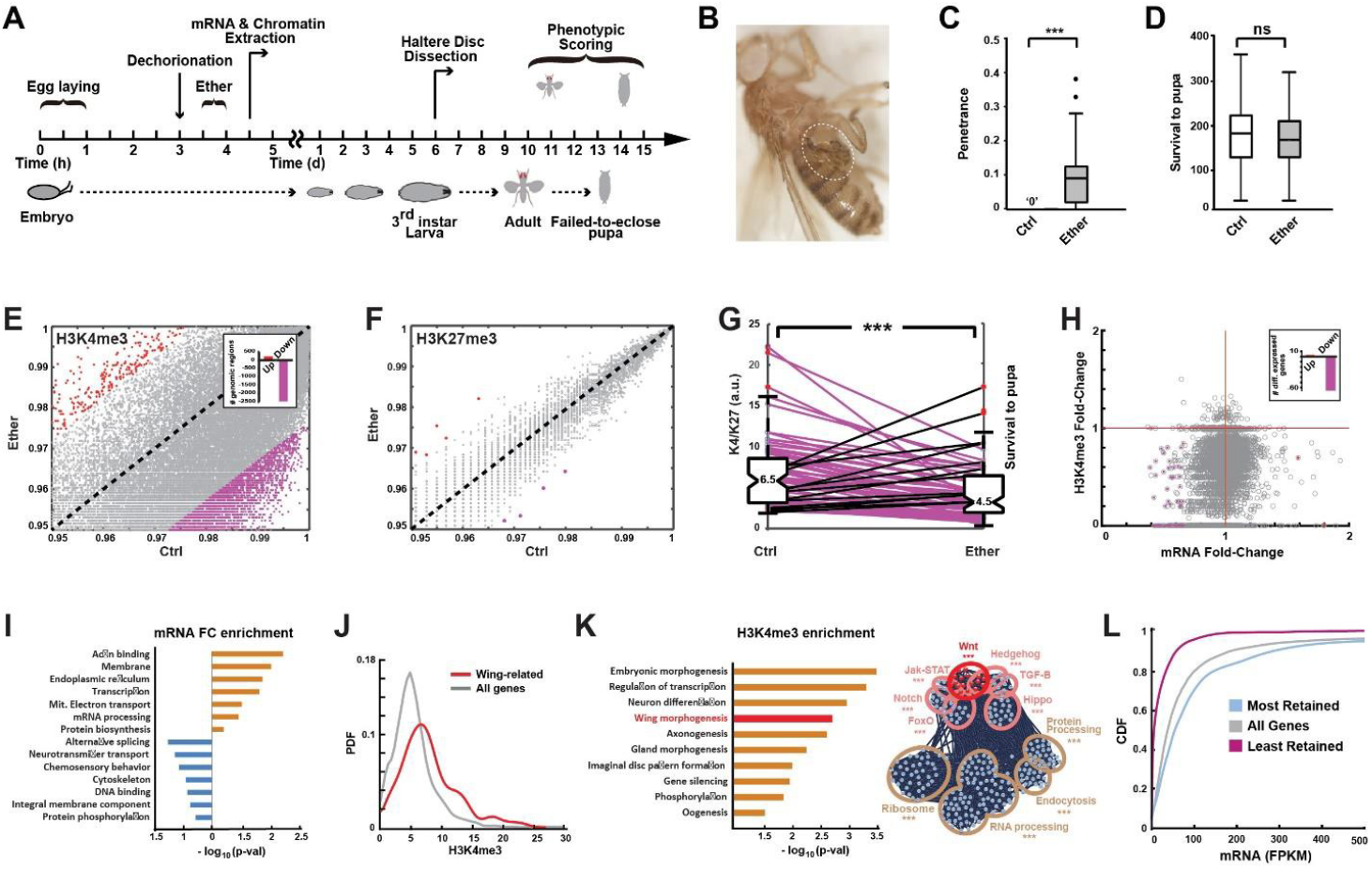
Ether suppresses H3K4 tri-methylation in exposed embryos but wing loci retain high levels. **(A)** Flowchart of experimental procedures and measurements. **(B)** Representative image of severe transformation in ether-exposed flies (*yw* line). **(C)** Fraction of individuals exhibiting bithorax phenocopies (penetrance) with and without ether exposure. Displayed fraction takes into account phenotypic adults as well as abnormal pupae that failed to eclose. **(D)** Survival to the pupal stage. **(E)** Percentile-normalized numbers of H3K4me3 reads per 100bp in ether-exposed (‘Ether’) vs. non-exposed *yw* embryos (‘Ctrl’). Red and purple dots correspond, respectively, to changes above and below two-standard deviations from the mean. Inset: no. of regions with methylation levels above and below 2 standard deviations from the mean. **(F)** Same as (E) for normalized numbers of H3K27me3 reads per 1000bp. **(G)** H3K4me3/H3K27me3 ratio per gene in ether-exposed vs. non-exposed embryos. *** p < 1E-19, Wilcoxon signed rank test. **(H)** mRNA fold-change (Ether/Ctrl) vs. H3K4me3 fold-change (Ether/Ctrl). Inset: numbers of differentially expressed genes (fold-change > 1.5 and p <0.05), based on 3 biological replicates. **(I)** Analysis of GO enrichment of transcripts that are up-regulated (red) and down-regulated (blue) 1hr after exposure vs. no exposure. Based on ‘DAVID’ online tool. **(J)** Computed probability density of H3K4me3 levels, corresponding to ‘wing development’ genes (red) and all genes (grey) of ether-exposed embryos. *** p < 1E-6. **(K) Left:** Functional enrichments of GO terms in genomic loci exhibiting the highest retention of H3K4 trimethylation (top 10%) following exposure to ether. Based on ‘DAVID’ online tool. Gene-specific retention is defined by the ratio between H3K4me3 levels in Ether-exposed and non-exposed embryos. **Right:** STRING Network analysis of genes with the highest retention of H3K4me3. Significance levels are indicated in Supplementary Table S1. **(L)** Cumulative distributions of mRNA (FPKM normalization), shown for all genes with a detectable level of H3K4me3 (grey) as well as for genes with 10% highest and lowest retention of H3K4me3 (blue and purple, respectively) *** p < 1E-6.

### Ether vapor causes disruptions in protein structure and reduces the level of trx protein

Since H3K4 tri-methylation in embryos is mediated primarily by trithorax function, the overwhelming decrease in H3K4me3 in ether-exposed embryos may be due to a negative impact on trx. This was consistent with decreased level of trx protein in exposed vs. non-exposed embryos (Fig. 2A). We then sought to investigate how the exposure to ether leads to reduction in trx function. Since ether is an organic solvent, we suspected that ether vapor has a denaturing impact on biomolecules, which may lead to rapid disruption of trx function due to proteotoxic stress. To investigate this possibility, we first analyzed the effect of ether on the integrity of the egg and its protein content. Inspection of embryos that were exposed to ether for 2h revealed gross increase in egg clarity (Fig. 2B), suggesting an increase in egg permeability and lipid precipitation. This was confirmed by elevated egg permeability to the nucleic-acid stain, Acridine Orange^19^ following exposure to ether vapor (Fig. 2C). To determine if the increased permeability (and/or vapor penetrance) affect the integrity of proteins in the embryo, we analyzed the total embryonic proteome by (Far-UV) circular dichroism (CD)^20,21^. Comparison of extracts from (*in-vivo*) exposed vs. non-exposed embryos revealed significant decrease in the degree of light polarization produced by proteins in the sample, suggesting a global impact on protein secondary structures, specifically α-helices (Fig. 2D). Staining with the protein conformation-sensitive probe, 8-anilino-1 naphthalene sulfonate (ANS)^22^, revealed concordant differences between extracts from exposed vs. non-exposed embryos. Notably, the direction of change in response to ether was the same as in heat denaturation (Fig. 2E). Independent analysis of the effect of ether on fluorescence intensity in live, Histone-RFP tagged (His2Av-mRFP1) embryos revealed a significant reduction of RFP intensity in ether-exposed vs control embryos (Fig. 2F). The reduction in His2Av-RFP fluorescence was observed without a change in His2Av mRNA (Supplementary Data S1), indicating post-transcriptional impact of ether on the His2Av-RFP protein. These findings indicate that the ether vapor compromises the integrity of the eggshell and causes disruptions of protein structure and/or function. This could potentially reduce the trx protein level by enhanced degradation of denatured trx and/or by an indirect impact of the proteotoxic stress response.

**Figure 2:**
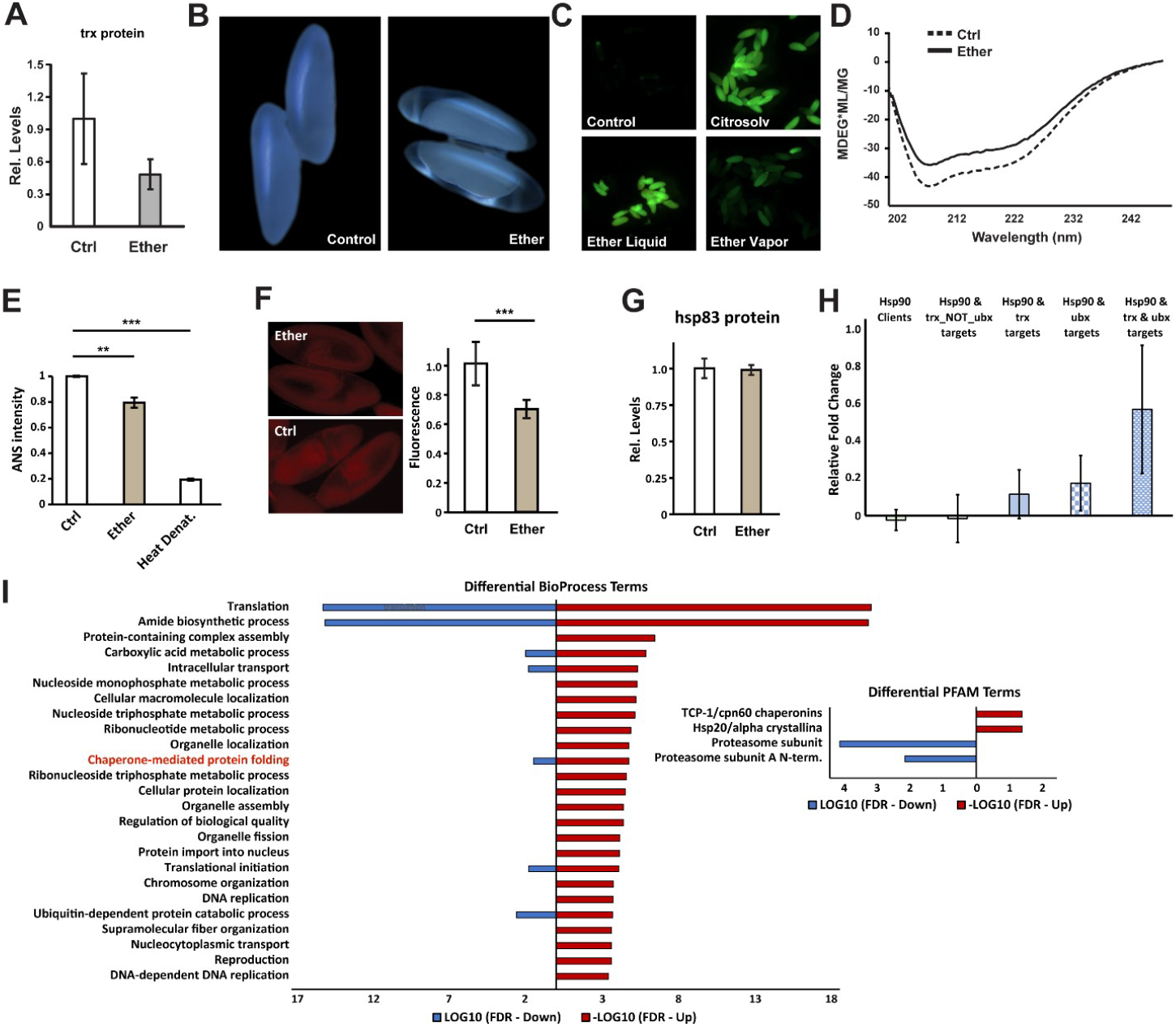
Ether induces proteotoxic stress and promotes differential chaperons’ activity of HSP90 and trx degradation. **(A)** Levels of the trx protein 1hr after exposure vs. no exposure. Mean ±SE, n=3. **(B)** Representative images of Acridine Orange-stained *yw* embryos for the following cases: no treatment (up left), 5min immersion in Citrasolv® solution (up right,^25^), immersion in ether liquid for 5 min (bottom left) and exposure to ether vapor for 1.5hr (Bottom right). **(C)** Circular dichroic (Far-UV) spectra of proteins extracted from yw embryos of exposure or no exposure to ether vapor (solid and dotted lines, respectively). Displayed spectra correspond to the average of 3 independent measurements. p < 0.05, Student’s t-test. **(D)** Fluorescence intensity of embryonic lysates (*yw* line) stained with 8-anilino-1-naphthalene sulfonate (ANS), a fluorescent probe for protein conformational changes. Shown for untreated embryos (Ctrl) and embryos that were exposed to ether vapor (Ether) or heat denaturation at 80° C. Mean intensity ± SE, based on 3 biological replicates. **p < 0.01, ***p < 0.001, One-way ANOVA following Dunnett’s test. **(E, F)** Representative images (E) and average fluorescence intensity in His2Av-mRFP1 tagged embryos (F), with and without exposure to ether vapor. Mean intensity per embryo ± SE, n=32 (Ctrl), n=29 (Ether). ***p < 1E-4, Mann-Whitney test. **(G)** Protein levels of the drosophila Hsp90 (hsp83) shortly after the exposure to ether. Based on proteomics analysis applied to exposed and non-exposed embryos 1hr after exposure. n=3. **(H)** Same as (F) for the indicated sub-groups of Hsp90 clients. **(I)** GO enrichment analysis of proteins that were increased (red) and decreased in response to ether, as determined by the proteomics profiles 1hr after exposure vs. no exposure. ** p < 0.01, *** p < 0.001.

### Ether exposure alters the deployment of Hsp90 clients

The proteotoxic stress caused by exposure to ether is expected to increase the workload on protein chaperones, such as Hsp90, potentially altering their target deployment^23,24^. Taken together with the reduced level of trx protein (Fig. 2A) and the reported dependence of trx function on Hsp90^11^, the altered deployment of Hsp90 (drosophila hsp83) might be responsible for the widespread suppression of H3K4 tri-methylation and may also account the induction of bithorax transformations in response to heat^12,13^. To investigate causal involvement of Hsp90 in bithorax induction, we analyzed the protein levels of Hsp90 and its targets. Proteomics analysis of whole embryos ∼1hr after exposure vs. no exposure, revealed that Hsp90 protein level was unaffected (Fig. 2G), but the levels of its wing-related clients were differentially increased by the exposure. This includes higher protein levels of Hsp90 clients that are also targets of Ubx and even more pronounced increase in clients that are jointly targeted by Ubx and trx (Fig. 2H). This contrasted with lack of change in Hsp90 clients that are targets of trx but not Ubx. Gene ontology analysis revealed significant enrichment of chaperones within subset of differentially elevated proteins and enrichment of proteosome genes in the subset of decreased proteins (Fig. 2I). The response to ether is therefore accompanied by differential increase of Hsp90 clients that are likely to favor bithorax transformation.

### Bithorax phenocopies are aggravated by joint reduction in *trx* and *Hsp90* function

To investigate potential involvements of trx and Hsp90 in the induction of bithorax phenocopies, we compared the outcomes of exposure on the background of *trx* and *Hsp90* mutants vs. wild-type background. The induced bithorax transformation in temperature sensitive *trx* stocks, *trx*^1-/+^ and *trx*^1-/-^, increased with mutation dosage and over 20% of *trx*^1-/-^ homozygotes exhibited spontaneous transformation without exposure to ether (Fig. 3A). Significant increase in the penetrance of bithorax phenocopies was also noted in *Hsp90* heterozygotes (*Hsp83*^*e6A*^) vs. wild-type control (Fig. 3B). Similar aggravation was observed in ether-exposed wildtype embryos that were pre-treated with the Hsp90 inhibitor, Geldanamycin, immediately after egg deposition (Fig. 3C). Analysis of *Hsp83*^*e6A*^/*trx*^1^ double heterozygotes showed that in contrast to *trx* and *Hsp90* single mutants, joint reduction in gene dosage of *Hsp90* and *trx* can cause spontaneous bithorax transformations in non-exposed embryos (Fig. 3D). Combined with the expected dependence of *trx* on the *Hsp90* function^11^, it suggests that the increase in bithorax reaction in embryos with reduced *Hsp90* function is due to excess reduction of *trx* function.

**Figure 3:**
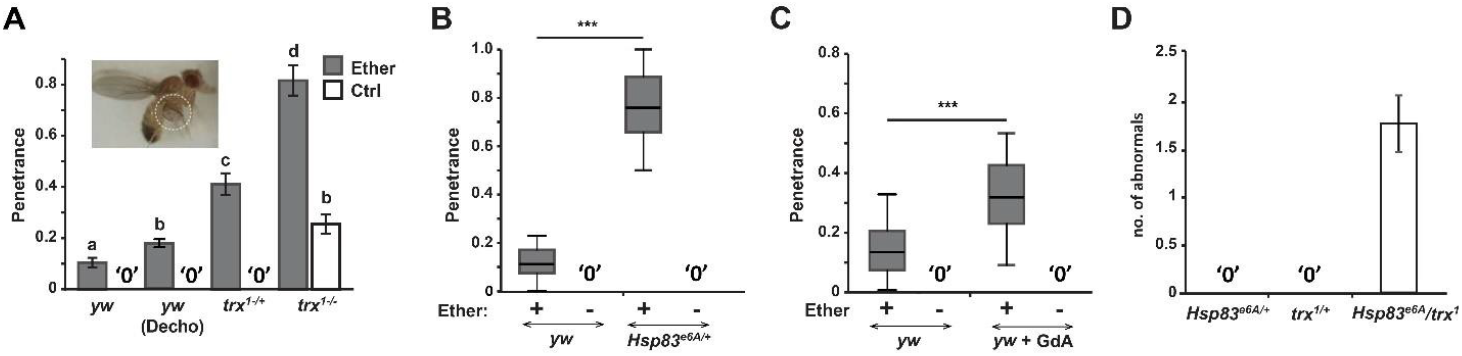
Bithorax epistasis between *Hsp90* and *trx* loss-of-functions. **(A)** Penetrance of bithorax phenocopies in ether-exposed and non-exposed adults from *trx*^1+/-^, *trx*^1-/-^ stocks vs. wild-type (*yw*), with and without egg dechorionation prior to exposure (‘Decho’). Mean penetrance ± SE. *a, b, c, d* denote significant difference (p < 0.05) compared with the respective control (Supplementary Table S2). Inset: Representative example of induced phenocopies in the *trx*^1^ line. **(B)** Penetrance of bithorax phenocopies in ether-exposed and non-exposed *Hsp83*^*e6A*^ stock vs. wild-type control (*yw*). **(C)** Sama as (B) on *yw* background, with and without egg treatment with the Hsp90 inhibitor, Geldanamycin (*yw* + GdA), prior to ether exposure. Mean penetrance ± SE, based on 3 biological replicates. *p < 0.05, **p < 0.01, Student’s t-test. **(D)** Average no. of *Hsp83*^e6A^/*trx*^1^, *trx*^1-/-^ and *Hsp83*^e6A^ flies exhibiting spontaneous phenocopies. Mean ± SE in. Based on 7 biological replicates.

### Ether exposure up-regulates wing-related *Ubx-trx* targets in the haltere

The enhanced bithorax reaction in *trx* mutants suggests that the ether-exposed decrease in the (*trx*-mediated) H3K4 tri-methylations is causally linked to the induction of bithorax phenocopies. To seek additional support, we first analyzed the effect of exposure on gene expression in haltere discs of 3rd instar larvae, with and without *trx* mutations (Supplementary Fig. S3A, Supplemental Spreadsheet S2). RNA-seq analysis of dissected discs revealed up-regulation of wing morphogenesis genes that increases with mutant dosage (Fig. 4A). This includes differential increase in the transcription of wingless (*wg*), the master regulator of wing development (Fig. 4B). Analysis of mRNA levels of *trx, Ubx* and their gene targets (*Ubx* itself is a trx target that specifies the haltere fate by repressing the transcription of wing genes^10,26– 28^). While the mRNA levels of *trx* and *Ubx* were largely unaffected by the exposure (Supplementary Fig. S3B-D; Supplemental Spreadsheet S2), the levels of *Ubx* targets were significantly up-regulated (Fig. 4C,D). The effects on targets of trx, on the other hand, depended on whether they are also targeted by *Ubx*; while joint targets of *trx* and *Ubx* were significantly up-regulated (Fig. 4C,D), targets of *trx* alone were strongly down-regulated (Fig. 4C,D). The relevance of these changes to induction of wing genes in the haltere was revealed by preferential up-regulation of wing genes that are targeted by *Ubx*, but not wing genes that are targeted only by *trx* (Fig. 4E,F). Since *Ubx* is itself a target of *trx* and a transcriptional repressor of wing genes in the haltere^24^, the up-regulation of joint *Ubx* and *trx* targets is consistent with the induction of wing phenotypes.

**Figure 4:**
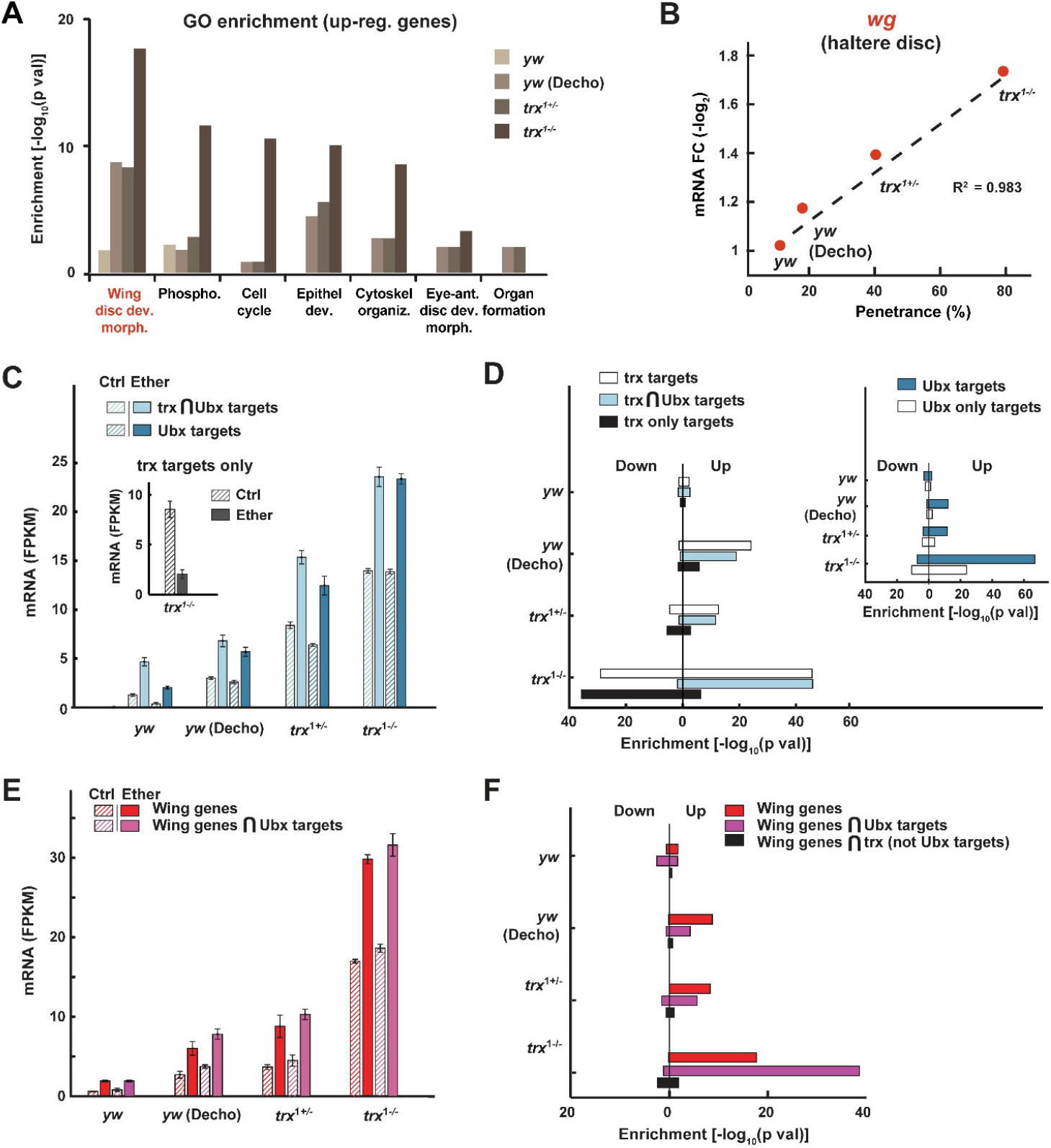
Bithorax induction is manifested by up-regulation of wing genes and *Ubx-trx* targets in the haltere. **(A)** Functional enrichments of GO terms in genes that were significantly up-regulated by ether in haltere discs of 3rd instar larvae from *trx*^1-/+^, *trx*^1-/-^ stocks and wild-type (yw) stock, with and without egg dechorionation prior to exposure (‘Decho’). Based on David online tool. **(B)** Ether-induced mRNA fold-change of wingless in the larval haltere disc. Mean fold-change (ether vs. control) for the cases in (A). **(C)** Median levels of mRNA ± SE for the indicated subsets of genes. n=3. Ether effect: p < 0.001, Genotype effect: p < 0.001, Ether-Genotype interaction: p < 0.001, Two-way ANOVA (full set of ANOVA p-values provided in Supplementary Table S3). **(D)** Enrichment of *trx* and *Ubx* targets (inset) within up- and down-regulated genes in haltere discs of 3rd instar larvae (fold-change > 1.5 and p <0.05), Fisher exact test. Shown for the cases in (A). **(E, F)** Same as (C, D) for wing development genes and their intersections with *Ubx* targets or with *trx* targets that are not *Ubx* targets. Ether effect: p < 0.001, Genotype effect: p < 0.001, Ether-Genotype interaction: p < 0.001, Two-way ANOVA (full set of ANOVA p-values provided in Supplementary Table S4).

### Upregulation of wing genes in the haltere correlates with H3K4me3 levels at the time of exposure

While the induction of wing phenocopies might be caused by the up-regulation of joint targets of *Ubx* and *trx* in the haltere, it is not clear how this up-regulation is promoted by brief exposure to ether in early-stage embryos. To investigate if the induced pre-disposition toward wing could be specified by the levels of H3K4 tri-methylation in the early embryo, we tested if the differential transcription of wing-related genes in the haltere disc correlates with the levels of K4 tri-methylation of these genes shortly after exposure. We found that genes with high and low H3K4me3 levels (top and bottom 10%, respectively) were expressed at high and low levels in the haltere (Fig. 5A). Reciprocal analysis of genes that are up-regulated in the haltere showed that at the time of exposure, these genes were more highly K4 tri-methylated compared with the bulk of up-regulated genes (Fig. 5B, C; Supplementary Fig. S4A). Preferential K4 tri-methylation at the time of exposure was even more pronounced in those wing-related genes that were found to be up-regulated in the haltere (Fig. 5C). Linkage between early H3K4 tri-methylation and later expression of wing genes in the haltere was further supported by differential methylation of specific subsets of *trx* targets (Fig. 5D). In particular, early-stage H3K4 tri-methylation of joint targets of *trx* and *Ubx* was significantly shifted towards higher levels (p < 1E-5), while H3K4 tri-methylation of *trx* targets that are not shared with *Ubx*, was shifted toward lower levels (p < 1E-5). These differences in H3K4me3 were consistent with matching (positive and negative) transcriptional fold-changes in the haltere discs (insets to Fig. 5D; Supplementary Fig. S4B, C). Taken together, these findings implicate wing gene related differences in H3K4 tri-methylation shortly after exposure with later-stage induction of haltere-to-wing transformations.

**Figure 5:**
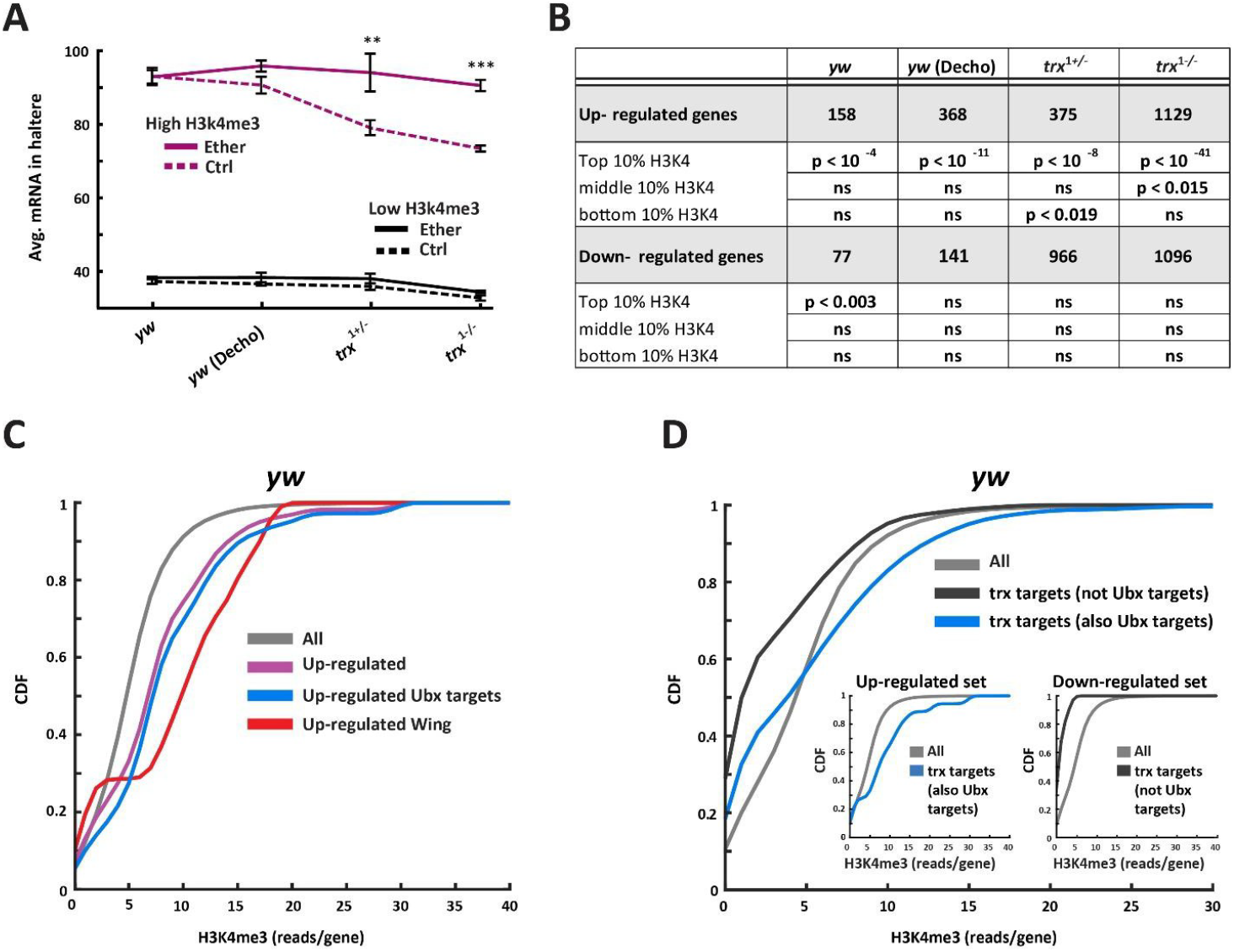
Up-regulation of wing genes in the haltere correlates with higher levels of H3K4me3 shortly after exposure. **(A)** Transcriptional levels in the haltere (3rd instar larvae) for genes with the highest (top 10%, purple) and lowest (bottom 10%, black) levels of H3K4me3 shortly after exposure to ether (4.5hr AED). Shown for yw (with and without dechorionation), *trx*^1-/+^ and *trx*^1-/-^. Mean FPKM ± SE, n=3. **p < 0.01, ***p < 0.001, Two-way ANOVA followed by Tukey HSD test. **(B)** Significant overlap between H3K4me3 levels in exposed embryos (yw line, 4.5hr AED) and differential expression (ether vs. control) in the haltere of disc of 3rd instar larvae. Shown for the cases in (A). Hypergeometric test. **(C)** Cumulative distributions of H3K4me3 levels, shown for up-regulated genes (purple), up-regulated Ubx targets (blue), up-regulated wing development genes (red) and all genes (gray) **(D)** Cumulative distributions of H3K4me3 for joint targets of *trx* and *Ubx* (blue), targets of *trx* that are not shared with *Ubx* (black) and all genes with detectable methylation (gray). Inset: Distributions of early levels of H3K4me3 (4.5hr AED) shown for the indicated subsets of targets that are up- and down-regulated in the haltere disc (left and right, respectively).

## Discussion

Fossil record evidence suggests that in contrast to flies, the common ancestor of all winged insects had two pairs of large membranous flight wings, located on the second and third thoracic segments^29^. The hindwings in different orders of insects evolved into organs with altered functions, such as the gyroscopic haltere in Drosophila. These alterations appear to have emerged as co-options of a wing program, as evidenced by reversals to a double pair of wings on the background of mutations in genes of the bithorax gene complex, such as *Ubx* and *trx*^10,30^. trx is the methylase responsible for histone H3K4 tri-methylation which contributes to maintenance of active state of expression of many genes, including the homeotic genes that specify segment identities in the Drosophila embryo (e.g. the Antennapedia and Bithorax genes)^31–33^. Ubx, in turn, is a trx target and a master regulator of haltere development that specifies haltere fates in Drosophila, by repressing the transcription of multiple wing genes in the third thoracic segment^34,3^. Loss of function mutations in these genes is therefore consistent with spontaneous haltere-to-wing transformations. In a seminal work on gene-environment interactions, Waddington demonstrated that haltere-to-wing transformations can also be induced by embryonic exposure of wild-type embryos to ether vapor^6^. Similar induction was demonstrated by exposure to heat at the same sensitivity time window^3–6,12,13,34^, but the mechanistic basis of induction has remained unknown for over 60 years. Our results show that brief exposure to ether at the time of cellularization alleviates the suppression of Ubx-dependent wing regulators in the larval haltere disc. This “reprogramming” of the larval disc is mirrored by relatively high H3K4 trimethylation of wing genes at the time of exposure. Preferentially higher H3K4me3 levels of these genes is, in turn, expected to assist in maintaining their active state of transcription over time^17^, consistent with the haltere-to-wing predisposition.

While investigating how early exposure to ether creates this predisposition, we discovered that ether vapor dissolves the egg shell and compromises protein integrity in the embryo. This increases the demand for Hsp90, a central chaperone that assists the folding of a wide range of proteins (‘clients’) during normal development and especially under stress. The chaperone activity of Hsp90 is required to support diverse functions, including transcriptional regulation, chromatin remodeling and phenotypic buffering of genetic and epigenetic variations^35–47^. Since Hsp90 has also been implicated in supporting trx function in Drosophila (and the methyltransferase function of SMYD3 in mammals)^11,48^, we suspected that the proteotoxic stress in ether-exposed embryos compromises trx function by altering the deployment of Hsp90. This was supported by several lines of evidence, including enhanced induction of bithorax phenocopies on the background of genetic or chemical reduction in Hsp90 and epistasis between reduced functions of *Hsp90* and *trx*. These findings implicate the environmental impact on chaperone function with altered pattern of epigenetic memory that predisposes the haltere segment for reversal toward wing.

By integrating our findings with evidence from previous studies, we propose a model that can account for bithorax-like transformations in response to early embryonic exposure to both ether and heat (Fig. 6; Supplementary discussion). Exposure at around the time of cellularization creates a proteomic stress followed by redeployment of the *Drosophila* Hsp90 (Hsp83) towards misfolded proteins. The reduced availability of Hsp90 for trx at a stage in which trx function is particularly required (Fig. 6,^49,50^) interferes with the setup of H3K4 tri-methylations. Genes with relatively high levels of H3K4me3 following the exposure (e.g. wing development genes that are targeted by Ubx) are expected to be pre-disposed for sustained expression at a later stage of development. This pre-disposition is most clearly manifested by alleviated suppression of Ubx targets in the third thoracic segment (whose identity is specified by Ubx), which is in turn, consistent with induction of wing phenotypes in the haltere.

**Figure 6:**
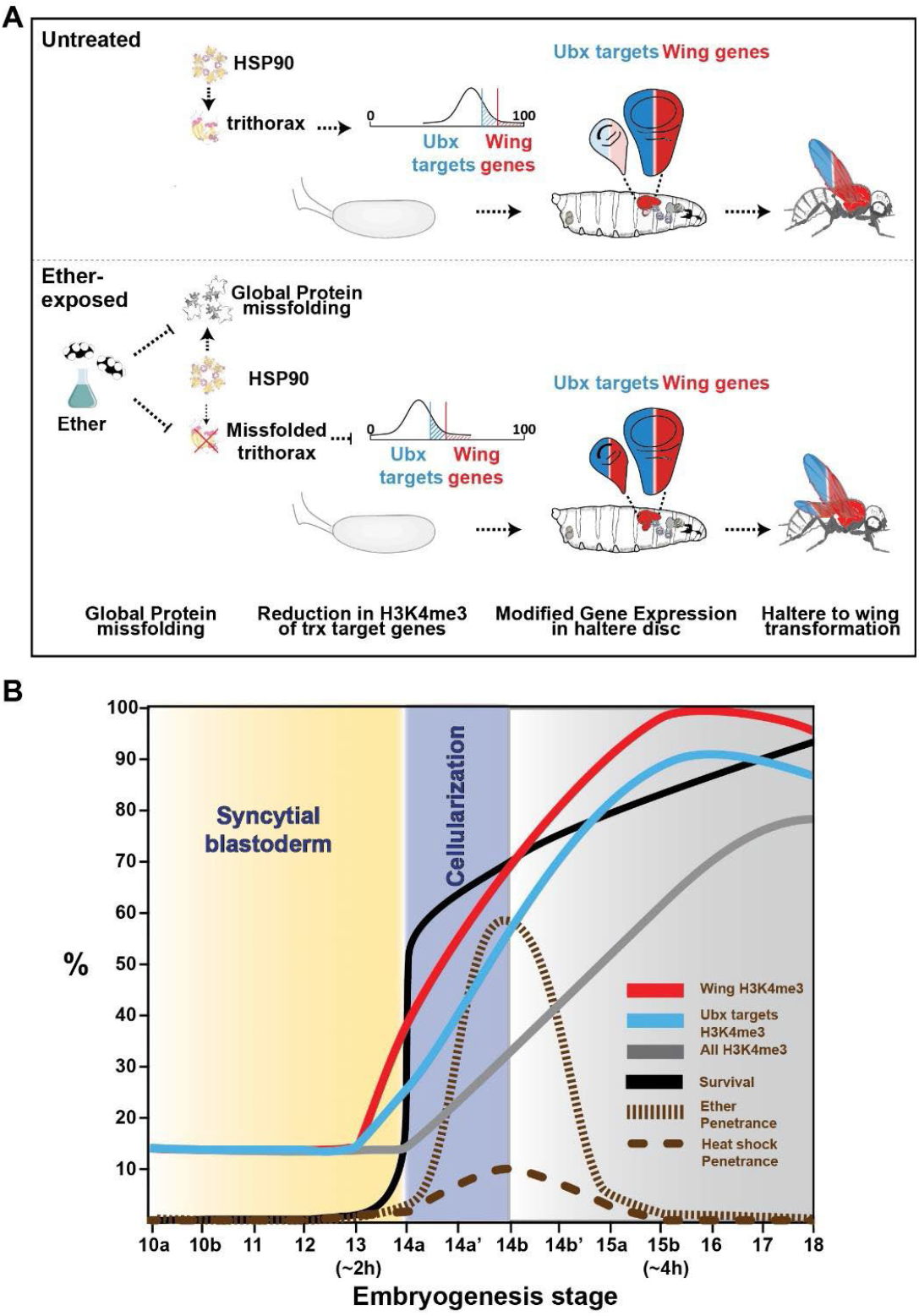
Hypothetical model of induced bithorax phenocopies. **(A)** Redeployment of Hsp90 towards misfolded proteins interferes with its contribution to trx function and compromising the establishment of H3K4me3 marks. If the exposure occurs shortly after the onset of H3K4 methylation, the patterns of H3K4me3 are not yet fully established. Genes that are differentially expressed prior this exposure are more highly k4 trimethylated and preferentially retain their H3K4me3 marks after the exposure. This set of genes is highly enriched with wing development genes that are normally repressed by Ubx expression in the haltere. The preferentially higher levels of H3K4me3 of these genes leads to more effective stabilization of their active state of transcription, leading to higher relative expression during later stages of development. The up-regulation of wing genes that are Ubx targets promotes haltere-to-wing transformation in the haltere. **(B)** Consistency of the model with multiple lines of independent evidence. The lethality of exposure during most the syncytial period^3,4,13^ is in line with increased permeability of ether vapor into the pre-cellularization embryo, which is expected to significantly aggravate the disruption of protein integrity in the egg cytoplasm. Enhancement of bithorax induction under exposure between stages 14 and 15 may be explained by the gradual priming of H3K4me3 in Ubx targets and wing development genes (modENCODE data^51^). The reduction in the efficacy of bithorax induction and the accompanying increase in survival to exposure after stage 15a^3,4,13^, are consistent, respectively, with substantial establishment of H3K4me3 marks at this stage^51^. The same intermediate ‘window of opportunity’ for bithorax induction (between stages 14 and 15) is also expected in the case of exposure to heat, thus accounting for the similarities in the effective stages and morphogenetic outcomes. The nomenclature of the developmental stages corresponds to that described by^13^.

Altogether, these findings portray a causal chain of events, connecting environmental disruption of protein integrity at onset of histone methylations with modified epigenetic patterning that supports morphogenetic shift towards an ancestral-like body plan. The increased homeotic sensitivity to reduction of Hsp90 and trx functions, likely applies to additional inductions of homeotic transformations^10,43^ and can be utilized for fate manipulation and/or regeneration at the levels of tissues and organs.

## Supporting information

Materials and Methods, References, Figs. S1 to S4, Tables S1 to S6, Captions for Data S1 to S2

## Acknowledgements

We thank Drs. Moshe Goldsmith, Maor Knafo and Yulia Gnainsky for helpful discussions. This work was supported by the Sir John Templeton Foundation (grant numbers: 40663 and 764 61122).

## Supplementary discussion

In addition to providing explanation for the induction of bithorax phenocopies by ether (or heat), the chain of events identified in this work can also account for the lethality of exposure at an earlier stage and the lack of responsiveness to exposure a few hours later (Fig. 6A). Previous work showed that the induction of bithorax phenocopies by either ether or heat shock is largely confined to a time window which opens shortly before cellularization and ends at around the stage of partial invagination of the anterior and posterior midgut^3,4,13^. Exposure during syncytial stages (<2hr AED) leads to complete embryonic lethality, while exposure after furrow formation (>4hr AED) is no longer capable of inducing a phenocopy. In between, the survival increases as a function of the onset of exposure, while the penetrance increases to a peak at around the end of cellularization and gradually decreases at later onsets of exposure (Fig. 6A). This phenomenology is fully consistent with the hypothesized involvement of Hsp90 and *trx* functions; early embryonic stages are characterized by rapid divisions of nuclei within a large cytoplasmic compartment. This cytoplasm is initially loaded with very high levels of maternal transcripts of Hsp90 (modENCODE data^52^) which contributes to protein folding and functional integrity in this large compartment. This function of Hsp90 may be particularly critical for maintaining the cytoplasmic protein gradients that specify the anterior-posterior and lateral-ventral axes^53^. Since the interruption of these gradients is lethal^54^, a sufficient disruption of protein integrity can account for the lethality of exposure at that stage (Fig. 6A). This was indeed supported by the two categories of defects that have been observed in the case of early exposure^3^, namely: (i) failure to form a blastoderm, resulting in an undifferentiated-like mass with no recognizable structures and (ii) emergence of anterior, posterior or segmentation defects that are eventually followed by failure to hatch. The abnormalities were also more pronounced in embryos that were exposed at progressively earlier stages^3^.

The phenotypic responsiveness, in turn, exhibits non-monotonic stage dependence which is consistent with the requirement for *trx* function at that particular stage^49,50^ and is accounted for by the hypothesized model. The latter proposes that the pre-disposition for haltere-to-wing transformation depends on having sufficient levels of H3K4me3 preferentially in wing genes at around the time of exposure to ether. Analysis of published H3K4me3 time series data ^51^, shows that the early H3K4me3 levels in Ubx targets and wing development gene loci are indeed higher than the median for the entire set of methylated genes. The differential accumulation of H3K4me3 in these wing-related genes begins at the late stage of syncytial blastoderm with the onset of the bithorax sensitivity window (Fig. 6B). The establishment of a Trithorax-based epigenetic memory is thought to depend on a sufficiently long duration of *trx* association with the respective gene locus^55,56^. Stable propagation of the active chromatin states may therefore require a sufficient dosage of functional trx protein^4,11,50^. The ether (or heat)-mediated repression of *trx* function can then lead to reduced expression stability preferentially for genes that were not sufficiently marked by trithorax function prior or during the exposure. The initially higher levels of H3K4me3 at Ubx target and wing gene loci therefore contributes to relatively higher expression stability, which is fully consistent with their later tendency of being up-regulated compared to other genes. This effect may act in concert with the determination of imaginal disc precursors from ectodermal cells, which takes place soon after cellular blastoderm formation^57–61^.

According to the proposed model, the predisposition towards higher relative levels of wing development genes is caused by impairment in *trx* function. This bias is gradually reduced as the onset of exposure gets closer to sufficient H3K4me3 at these loci, thus reducing the efficacy of induction. This is consistent with the observation that the end of the bithorax responsiveness window coincides with the end of critical requirement for *trx* function^10,49,62^ as well as with a reported halt in accumulation of H3K4me3 across the genome (Fig. 6B) ^51^. Since re-deployment of Hsp90 and reduction in H3K4 tri-methylation are expected in ether and heat exposures, the proposed model also accounts for the striking overlap between the respective windows of responsiveness.

The hypothesized model therefore suggests the following axis of influence: Early exposure **→** reduced integrity of proteins **→** altered deployment or function of Hsp90 **→** compromised *trx* function **→** altered profile of H3K4me3 **→** predisposition for upregulation of Ubx targets and wing regulators **→** homeotic transformations. The increased homeotic sensitivity to reduction of Hsp90 and *trx* functions, likely applies to additional inductions of homeotic transformations^10,43^ and can be utilized for fate manipulation and/or regeneration at the levels of tissues and organs.

