## Supplementary material for "Waddington Revisited: Organ transformation by environmental disruption of epigenetic memory": Materials and Methods, References, Figs. S1 to S4, Tables S1 to S6, Captions for Data S1 to S2

**This file includes:**

Materials and Methods
References
Figs. S1 to S4
Tables S1 to S6
Captions for Data S1 to S2

**Other Supplementary Materials for this manuscript include the following:**

Data S1 to S2

### Materials and Methods

#### *Drosophila* stocks

*Drosophila* lines *trx*$^{1}$*/TM1*, *Hsp83*$^{e6A}$ were obtained from the Bloomington Stock Center. The *yw* stock was obtained from the laboratory of Prof. Eli Arama (Weizmann Institute of Science, Israel).

#### Generation of *trx*$^{1}$*/ TM6B, Tb* flies

We have replaced the third chromosome balancer of the original line *trx*$^{1}$*/TM1* with a Tm6b balancer carrying a larval marker (Tb) to have the ability to differentiate between homozygous and heterozygous larvae. We have restored the genetic background of the original line[1], excluding the Tm6b, using chromosomal markers.

#### Food preparation

Standard cornmeal food (Bloomington Stock Center recipe, http://flystocks.bio.indiana.edu/Fly_Work/media-recipes/molassesfood.htm).

#### Exposure and scoring of responses to ether

0-3 day old flies were reared on fly medium supplemented with dry yeast for 3 days under 12h light/dark cycle regime, temperature of 25°C and 70% humidity. About 10,000 adult flies were taken for 5 rounds of egg deposition, 1h each, in cages on a 10 cm agar plate supplemented with dry yeast. The first two rounds of egg deposition were discarded for embryo's developmental stage synchronizations. Dechorionation was performed 2 h later, using 3% sodium hypochlorite for 2.5 min.  Following 2.5-3.5h after oviposition[2], at the embryonic syncytial blastoderm stage, the eggs were exposed to ether. Eppendorf tube containing 1.5 ml of diethyl ether was placed in each glass bottle, allowing the ether vapor to diffuse and exposing the embryos to a fixed concentration of ether vapor in 25˚C for 30 min. Then, tubes containing the remaining ether liquid were removed and the bottles with the embryos were left to evaporate any ether remains. For embryonic RNA and ChIP analyses, 3:45-4:45 h old embryos were collected and flash freezed in liquid nitrogen. For haltere disc RNA analysis and phenocopy scoring, embryos were transferred to new clean bottles and allowed to hatch. We have dissected haltere discs from 3rd stage larvae developed from ether exposed and non-exposed embryos. Phenocopy scoring was done every day from the first day of eclosion till 5 days later. After the fifth day all unenclosed pupae were dissected and presence of phenocopy was determined. For *Hsp90* inhibition experiments, embryos were dechorionated immediately after oviposition and rocked in PBS supplemented with Geldanamycin (35 µM) for 1h[3].

#### Chromatin immunoprecipitation (ChIP) and ChIP-seq

Ether exposed and non-exposed embryos (3:45-4:45h old) were collected (0.1 mg per group). Embryos were crosslinked in 1 ml A1 buffer (60 mM KCl, 15 mM NaCl, 15 mM HEPES [pH 7.6], 4 mM MgCl2, 0.5% Triton X-100, 0.5 mM dithiothreitol (DTT), and complete EDTAfree protease inhibitor cocktail [Roche]), in the presence of 1.8% formaldehyde and homogenized at the same time in a douncer followed by incubation for 15 min at room temperature. Crosslinking was stopped by adding 225 mM glycine followed by incubation for 5 min. The homogenate was transferred to a 1 ml tube and centrifuged for 5 min, 4,000 x g at 4°C. The supernatant was discarded, and the nuclear pellet was washed three times in 3 ml A1 buffer and once in 3 ml of A2 buffer (140 mM NaCl, 15 mM HEPES [pH 7.6], 1 mM EDTA, 0.5mMEGTA, 1%Triton X-100, 0.5mMDTT, 0.1% sodium deoxycholate, and protease inhibitors) at 4°C. After the washes, nuclei were resuspended in A2 buffer in the presence of 0.1% SDS and 0.5% N-lauroylsarcosine and incubated for 30 min on a rotating wheel at 4°C. Chromatin was sonicated using a Bioruptor (Diagenode) for 15 min (settings 30s on, 30s off, high power). Sheared chromatin had an average length of 300 to 700 base pairs. After sonication and 10 min high-speed centrifugation, fragmented chromatin was recovered in the supernatant. Chromatin was precleared by addition of 50 µl of Protein A-Agarose (PA) suspension (Roche 11134515001) followed by overnight incubation at 4°C. PA was removed by centrifugation, antibodies at dilution 1:100 were added to the supernatant (a control in the presence of rabbit preserum [Mock IP] was performed at the same time), and samples were incubated for 4 hr at 4°C in a rotating wheel. PA (50 µl) was added, and incubation was continued overnight at 4°C. Antibody-protein complexes were collected by centrifugation at 4,000 rpm for 1 min, and the supernatants were discarded. Samples were washed four times in A3 (A2+ 0.05% SDS) buffer and twice in 1 mM EDTA, 10 mM Tris (pH 8) buffer (each wash, 5 min at 4C). Chromatin was eluted from PA in 250 µl of 10 mM EDTA, 1% SDS, 50 mM Tris (pH 8) at 65°C for 15 min, followed by centrifugation and recovery of the supernatant. The eluate was incubated overnight at 65°C to reverse crosslinks and treated with Proteinase K for 3 hr at 50°C. Sodium acetate (110 µM) was added to the samples, phenol-chloroform extracted, and ethanol precipitated in the presence of 20 µg glycogen. DNA was resuspended in 100 µl of water. Deep sequencing analysis of DNA was performed by Fasteris SA (Geneva, Switzerland) ChIP-seq library preparation was performed with illumina TruSeq ChIP kit. Adaptors were removed from raw FASTQ reads with cutadapt. The reads were then aligned to drosophila genome (UCSC dm3) with bowtie2; samtools and bedtools were used to convert resulting SAM files to the required downstream formats (bedraph etc.). We performed the analysis via the "Misha" R package[4]. The signal was smoothed via moving window averaging over 100bp followed by global percentile normalization. Next a 95% threshold was applied to separate signal from the background.

**RNA-seq library preparation and sequencing**

The cDNA libraries were prepared from poly-A mRNA following the manufacturer's instructions in the Illumina RNA sample preparation kit. In short, poly (T) oligo-attached magnetic beads were used to purify the poly(A)-containing mRNA molecules. The mRNA was fragmented into 200 to 500bp segments. RNA fragments were converted into cDNA using SuperScript II reverse transcriptase (Life Technology) and random hexamer primers. Adaptors were ligated to the cDNA fragments, followed by purification, PCR and additional purification. Deep sequencing measurement of RNA was performed in the Genomics Core Facility unit of the Technion Genome Center, Technion -Israel Institute of Technology (Haifa, Israel) using Illumina Genome Analyzer IIx (GA IIx). For sequencing, we used the following experimental kits and reagents: 1) Standard Cluster Generation Kit (#GD-103-4001, Illumina, San Diego, CA, USA) containing all reagents necessary to load the samples on to the flow cell and perform the bridge amplification, 2) Illumina Sequencing Kit v5 (TruSeq SBS Kit v5 GA (36-cycles), FC-104-5001) which contains the reagents for the sequencing runs, and 3) the GA IIx Sequencing Control Software version SCS 2.8, which was used to control the sequencer. Sequencing was based on 50bp single-end reads. mRNA was barcoded in the ligation step by Illumina standard multiplex adaptors. The multiplexed samples were sequenced on a single lane to yield between 2 and 8 million reads per sample.

**RNA-seq analysis**

Adaptors were removed from sequence reads using the cutadapt program[5]. Reads were mapped to the drosophila transcriptome (Ensembl version BDGP.25) using Bowtie2 and TopHat software[6], then Cufflinks and Cuffmerge[6] were applied to define a list of transcripts that are comparable between all samples. Differentially expressed transcripts including fold-change and statistics were identified by applying the DESeq R package[7] on the bowtie2 output.

GO enrichments were computed using the 'DAVID' online resource[8,9] with cutoffs for up/down regulation and FDR set to 1.5-fold and 0.05, respectively. Up- and down-regulated gene-sets were analyzed separately.

**Eggshell permeabilization**

Several hundred of *yw* adult flies were synchronized twice for 1 h and allowed to lay eggs for 1h on a 10 cm agar plate. Eggs were collected from the plate, washed in water, dechorionated and immersed in Citrasolv® (Citra Solv, Danbury, Connecticut)(1:10 dilution, 5 min) , diethyl ether (5 min) or were exposed to diethyl ether vapors for 1.5h in a closed bottle. Control embryos were left untreated in a closed bottle for the same period of time. Then the embryos were stained with Acridin orange dye for 5 minutes. Images were taken using a fluorescent stereoscope LEICA MZ16F equipped with Nikon digital sight DS-Fi1 camera.

**Circular dichroism and Fluorescence Spectroscopy**

CD spectra were recorded on a Chirascan spectropolarimeter (Applied Photophysics) calibrated with a solution of ammonium d-10-camphorsulfonate. Far-UV CD spectra were acquired using 1-mm path-length cuvettes, a step size of 0.5 nm, a bandwidth of 1 nm, and a time constant of 1 s. Protein concentration was 0.4 mg/mL.

Fluorescence emission (0.1-0.4 mg/mL protein, 25 °C; $\lambda ex$ = 380 nm, $\lambda em$ = 545 nm for unbound dye and $\lambda em$ = 470 nm for the dye-protein complex) were recorded with a Cytation5 (BioTek) plate reader fluorimeter in standard 96-well plate. In the experiments conducted in the presence of 8-Anilinonaphthalene-1-sulfonic acid (ANS) (1 mM; protein concentration, 0.1-0.4 mg/mL) the excitation wavelength was 380 nm. Before recording, the proteins were let to interact with the dye for 1 h. Following acquisition, both experiments (CD and fluorescence) were corrected for buffer (and dye, in the experiments involving ANS) contributions, averaged, and smoothed using sliding windows of 1.5 nm (far- and near-UV CD) or 3 nm (fluorescence).

**Quantification of RFP Fluorescence**

Several hundred of His3Av-mRFP1 adult flies were synchronized twice for 1 h and allowed to lay eggs for 1h on a 10 cm agar plate. Eggs were collected from the plate, washed in water, dechorionated and exposed to ether as previously described. Eggs were then transferred to MatTek Glass-Bottom Dishes that were pre-treated with embryo glue (3 M tape in Heptane) and covered in Halocarbon oil 700 (Sigma). The eggs were imaged using UPLSAPO 20 × numerical aperture: 0.75 objective of the confocal OLYMPUS FV1000 microscope with temperature-controlled chamber (set at 25 °C) and IX81 ZDC Motorized Stage. Image analysis was performed using ImageJ.

**Proteomics**

The samples were subjected to tryptic digestion using an S-trap. The resulting peptides were fractionated offline using high pH reversed phase chromatography, followed by online nanoflow liquid chromatography (nanoAcquity) coupled to high resolution, high mass accuracy mass spectrometry (Q Exactive HF). Each sample was analyzed on the instrument separately in a random order in discovery mode. Raw data was processed with MaxQuant v1.6.0.16. The data was searched with the Andromeda search engine against the uniprot drosophila melanogaster proteome database appended with common lab protein contaminants and the following modifications: Carbamidomethylation of C as fixed modification and oxidation of M and deamidation of NQ as variable ones. The LFQ (Label-Free Quantification) intensities were calculated and used for further calculations using Perseus v1.6.0.7. Decoy hits were filtered out, as well as proteins that were identified on the basis of a modified peptide only. The ratio of the LFQ intensities between the different samples was calculated and GO annotations added.

**Statistical analyses**

Statistical tests were performed using MATLAB (MathWorks) and R statistical program[10]. The significant difference between a subset group of genes to the entire population in their H3K4me3 or mRNA levels was numerically calculated using a bootstrap-based statistical test, as follows: this test was based on repeated cycles (1,000,000) of selecting genes at random from the total set of genes (same sample size as the subset group) and counting the fraction of times, in which the median methylation/expression in this random selection exceeds or fell behind the median level of the true subset. Significance was determined based on the percentage of iterations in which this analytical p-value was equal to, lower or higher than the median of the total set of genes. Analysis of enrichment of gene ontology annotations in sets of up- and down-regulated genes was done using the DAVID web tool with Benjamini correction for multiple hypothesis testing [8,9].

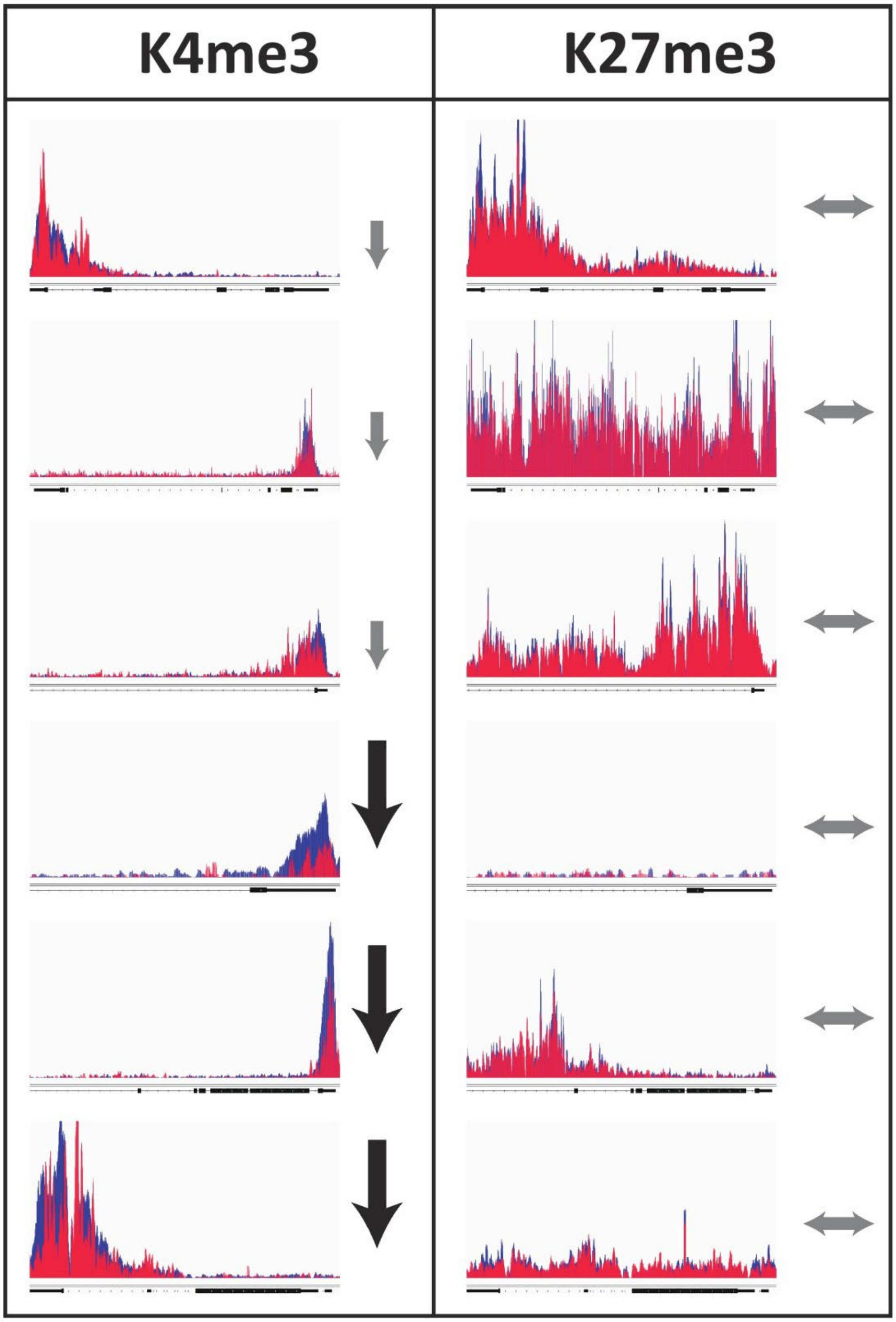


**Figure S1: Representative ChIP-seq profiles**. *wg*, *abd-A, tup, BicD, ssp3* and *tsh* in ether-exposed (red) and non-exposed embryos (blue). Left and right panels correspond to H3K4me3 and H3K27me3 marks, respectively. Vertical and horizontal arrows indicate, respectively, reduced number of reads and lack of change.

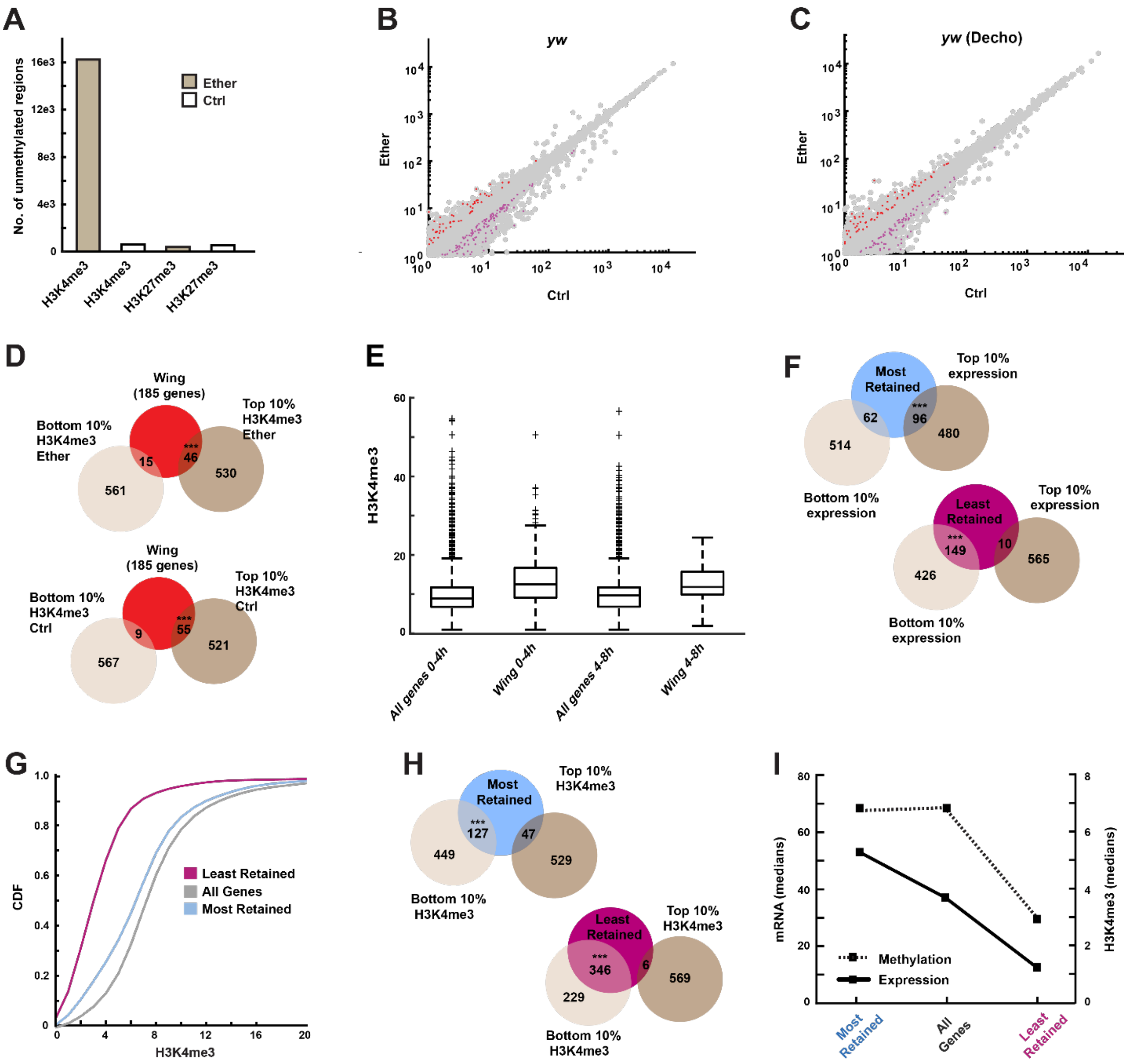


**Figure S2: Ether suppresses H3K4 tri-methylation, primarily in low-expressed loci**. **(A)** Normalized numbers of genomic regions with H3K4 and H3K27 tri-methylation (100bp and 1000bp long, respectively), measured for ether-exposed and control embryos (*yw*). **(B, C)** mRNA levels for ether-exposed and control embryos, with and without dechorionation (C and B, respectively). Differential expression (absolute fold-change > 1.5, p < 0.05, n = 3) is indicated by red and purple overlays. **(D)** Intersection between wing disc development genes and genes with highest and lowest H3K4me3 levels (top and bottom 10%) shortly after exposure to ether and without. *** p < 1E-8, *** p < 1E-14, respectively. hypergeometric test. **(E)** Box plots of H3K4me3 read counts corresponding to two time intervals of embryonic development (0-4hr and 4-8hr AED). Displayed for all genes and wing development genes ('Wing') based on compilation of ModEncode data [11]. **(F)** Intersection between genes with high preferential retention of H3K4me3 marks ('Most retained', blue) and genes with highest and lowest expression in control embryos. *** p < 1E-7. Same for intersection of 'Least retained' genes (purple). *** p < 1E-22. **(G)** Cumulative distributions of normalized H3K4me3 level per gene, shown for all genes with any detectable H3K4me3 (grey); and genes with 10% highest and lowest retention of H3K4me3 (blue and purple, respectively). **(H)** Intersection between genes with high preferential retention of H3K4me3 marks (‘Most retained’, blue) and genes with the highest and lowest H3K4me3 levels in control embryos (top and bottom 10%). *** $p < 1E-19$, hypergeometric test. Same for intersection of ‘Least retained’ genes (purple). *** $p < 1E-137$. **(I)** Median mRNA (solid line) and H3K4me3 levels (dashed line) for genes with high, medium and no preferential retention.

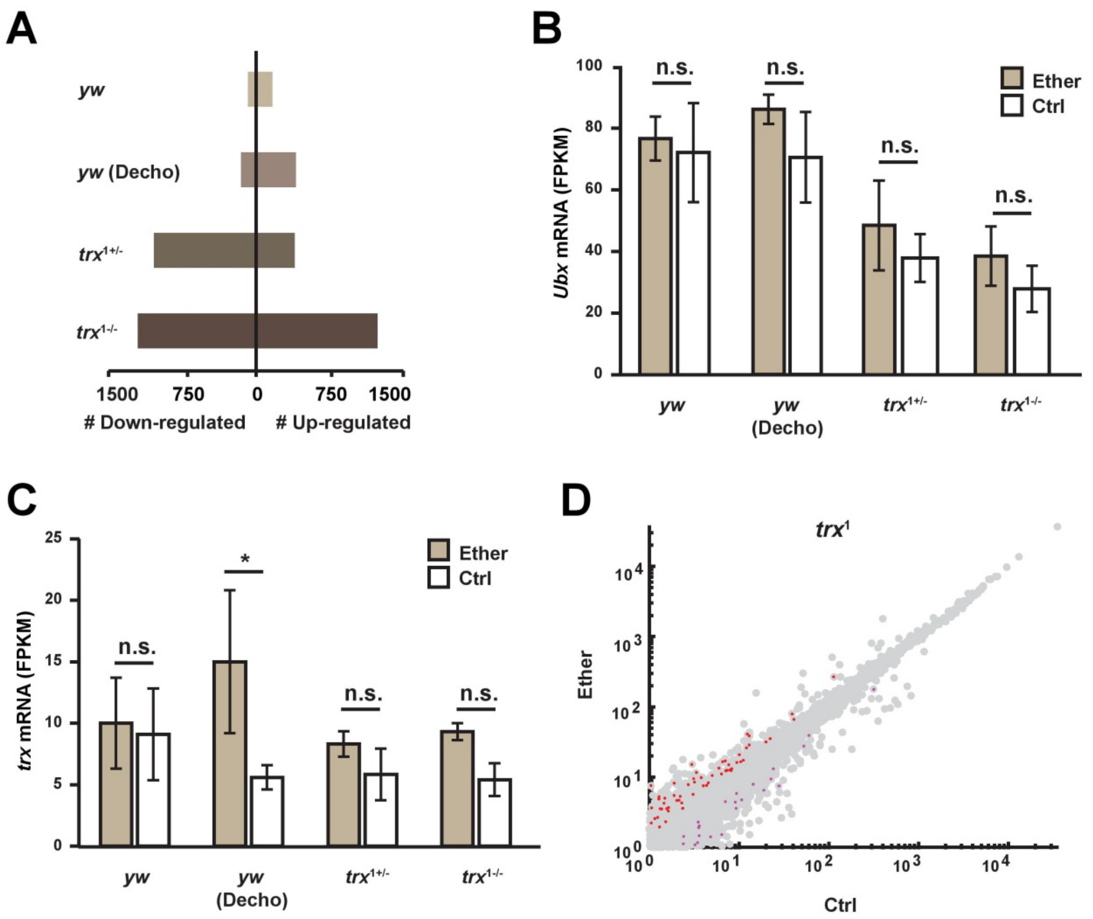


**Figure S3:** **(A)** Number of differentially expressed genes (absolute fold-change > 1.5, p < 0.05, n=3) in ether-exposed *yw*, *yw* decho, $trx^{1+/-}$ and $trx^{1-/-}$ haltere imaginal discs. **(B)** *Ubx* mRNA in ether-exposed *yw*, *yw* decho, $trx^{1+/-}$ and $trx^{1-/-}$ haltere imaginal discs Average ± SE, n=3. Two-way ANOVA following Tukey HSD test. **(C)** Same as (B) for *trx* mRNA. Two-way ANOVA following Tukey HSD test. **(D)** mRNA levels in $trx^1$ embryo shortly after exposure to ether vs. control. Differential expression (absolute fold-change > 1.5, p < 0.05, n=3) is indicated by red and purple overlays.

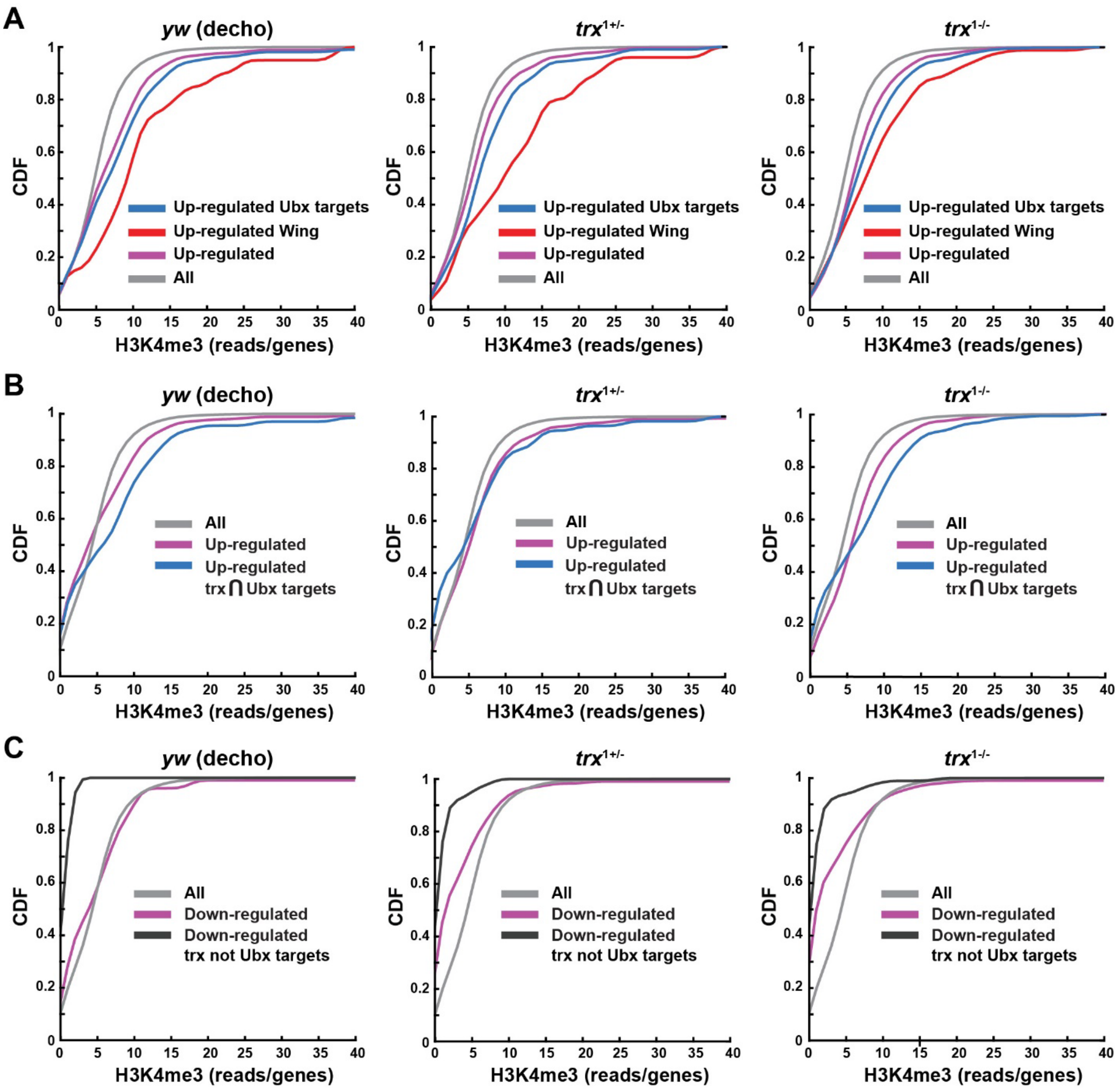


**Figure S4: Up-regulation of wing genes in the haltere correlates with high H3K4me3 at the time of exposure**. **(A)** Cumulative distributions (CDFs) of H3K4me3 levels shortly after ether exposure, shown for all genes with detectable H3K4me3 (grey), genes that are significantly up-regulated in the haltere disc (pink), up-regulated wing development genes and up-regulated Ubx targets (red and blue, respectively). Up-regulated gene sets in each panel are based on expression measurements in dechorionated *yw* flies (left), $trx^{1+/-}$ flies (center) and $trx^{-1-/-}$ (right). CDFs are based on the sum of normalized H3K4me3 reads for each gene. **(B)** Same as (A) with replacement of the last two groups with up-regulated genes that are joint targets of trx and Ubx (blue). **(C)** Same as (B) with replacement of the last two groups with significantly down-regulated genes in the haltere disc (pink) and down-regulated genes that are joint targets of trx and Ubx (black).

**Table S1. Gene network enrichment analysis of the genes with promoter regions within the highest H3K4me3 retention levels (top 10%)**.

| Gene network | Adjusted p-value |
|---|---|
| Spliceosome | 8.58E-32 |
| Ribosome | 2.81E-31 |
| Hippo signaling pathway - fly | 1.35E-28 |
| RNA degradation | 2.96E-24 |
| Ubiquitin mediated proteolysis | 9.23E-24 |
| Endocytosis | 5.66E-23 |
| RNA transport | 2.96E-20 |
| FoxO signaling pathway | 2.96E-20 |
| Wnt signaling pathway | 3.09E-19 |
| Protein processing in endoplasmic reticulum | 2.80E-15 |
| mRNA surveillance pathway | 1.31E-14 |
| Hedgehog signaling pathway | 6.61E-14 |
| TGF-beta signaling pathway | 1.04E-11 |
| mTOR signaling pathway | 2.12E-11 |
| Jak-STAT signaling pathway | 1.87E-08 |
| Notch signaling pathway | 4.31E-08 |
| Phagosome | 0.00348 |
| Regulation of autophagy | 0.0365 |

**Table S2. Post-hoc p-values for gene-environment interaction effects on penetrance**. Based on the fraction of pupae and adults presenting bithorax phenocopies in each bottle. Environmental conditions: Ether, Dechorionation ('Decho') and no treatment control. Genotypes: *yw*, $trx^{1+/-}$ and $trx^{1-/-}$. Paired Tukey HSD analysis applied to each of the indicated pairs.

| Group 1 | Group 2 | Adjusted p-value |
|---|---|---|
| $trx^{1-/-}$:(-)Ether | $trx^{1+/-}$:(-)Ether | 0 |
| *yw*:(-)Ether | $trx^{1+/-}$:(-)Ether | 1 |
| *yw* Decho:(-)Ether | $trx^{1+/-}$:(-)Ether | 1 |
| $trx^{1+/-}$:(-)Ether | $trx^{1+/-}$:(+)Ether | 0 |
| $trx^{1-/-}$:(+)Ether | $trx^{1+/-}$:(-)Ether | 0 |
| *yw*:(+)Ether | $trx^{1+/-}$:(-)Ether | 0.0000004 |
| *yw* Decho:(+)Ether | $trx^{1+/-}$:(-)Ether | 0 |
| *yw*:(-)Ether | $trx^{1-/-}$:(-)Ether | 0 |
| *yw* Decho:(-)Ether | $trx^{1-/-}$:(-)Ether | 0 |
| $trx^{1+/-}$:(+)Ether | $trx^{1-/-}$:(-)Ether | 0.048322 |
| $trx^{1-/-}$:(+)Ether | $trx^{1-/-}$:(-)Ether | 0 |
| *yw*:(+)Ether | $trx^{1-/-}$:(-)Ether | 0.0057505 |
| *yw* Decho:(+)Ether | $trx^{1-/-}$:(-)Ether | 0.6117278 |
| *yw* Decho:(-)Ether | *yw*:(-)Ether | 1 |
| $trx^{1+/-}$:(+)Ether | *yw*:(-)Ether | 0 |
| $trx^{1-/-}$:(+)Ether | *yw*:(-)Ether | 0 |
| *yw*:(+)Ether | *yw*:(-)Ether | 0.0000004 |
| *yw* Decho:(+)Ether | *yw*:(-)Ether | 0 |
| $trx^{1+/-}$:(+)Ether | *yw* Decho:(-)Ether | 0 |
| $trx^{1-/-}$:(+)Ether | *yw* Decho:(-)Ether | 0 |
| *yw*:(+)Ether | *yw* Decho:(-)Ether | 0.0000004 |
| *yw* Decho:(+)Ether | *yw* Decho:(-)Ether | 0 |
| $trx^{1-/-}$:(+)Ether | $trx^{1+/-}$:(+)Ether | 0.000002 |
| *yw*:(+)Ether | $trx^{1+/-}$:(+)Ether | 0 |
| *yw* Decho:(+)Ether | $trx^{1+/-}$:(+)Ether | 0.0000479 |
| *yw*:(+)Ether | $trx^{1-/-}$:(+)Ether | 0 |
| *yw* Decho:(+)Ether | $trx^{1-/-}$:(+)Ether | 0 |
| *yw* Decho:(+)Ether | *yw*:(+)Ether | 0.0418766 |

**Table S3: ANOVA p-values for the effects of the indicated factors on mean expression of subsets of trx and Ubx targets in the haltere disc of 3rd instar larvae**.

| Set of genes | Factor | p-value |
|---|---|---|
| Ubx targets | Genotype | 2.00E-16 |
| | Ether | 2.68E-11 |
| | Genotype:Ether | 7.79E-07 |
| trx & Ubx targets | Genotype | 6.34E-15 |
| | Ether | 1.08E-11 |
| | Genotype:Ether | 9.95E-05 |
| trx but not Ubx targets | Genotype | 2.28E-08 |
| | Ether | 1.33E-08 |
| | Genotype:Ether | 3.64E-05 |

.

**Table S4: ANOVA p-values for the effects of the indicated factors on mean expression of wing-related targets of Ubx in the haltere disc of 3rd instar larvae**.

| Set of genes | Factor | p-value |
|---|---|---|
| Wing-related genes | Genotype | 8.91E-16 |
| | Ether | 1.51E-09 |
| | Genotype:Ether | 8.51E-07 |
| Wing-related genes (also Ubx targets) | Genotype | 8.62E-16 |
| | Ether | 1.28E-09 |
| | Genotype:Ether | 1.57E-06 |

**Data S1. (separate file)**

**FPKM values in ether-exposed and non-exposed *yw*, *yw* decho, *trx*$^{1+/-}$ and *trx*$^{1-/-}$ embryos.**

**Data S2. (separate file)**

**FPKM values in ether-exposed and non-exposed *yw*, *yw* decho, *trx*$^{1+/-}$ and *trx*$^{1-/-}$ 3$^{rd}$ larvae haltere imaginal discs.**
